# Costs of Clock-Environment Misalignment in Individual Cyanobacterial Cells

**DOI:** 10.1101/045146

**Authors:** Guillaume Lambert, Justin Chew, Michael J. Rust

## Abstract

Circadian rhythms are endogenously generated daily oscillations in physiology found in all kingdoms of life. Experimental studies have shown that the fitness of *Synechococcus elongatus*, a photosynthetic microorganism, is severely affected in non-24h environments. However, it has been difficult to study the effects of clock-environment mismatch on cellular physiology because such measurements require the precise determination of both clock state and growth rates in the same cell. Here, we designed a microscopy platform that allows us to expose cyanobacterial cells to pulses of light and dark while quantitatively measuring their growth, division rate, and circadian clock state over many days. Our measurements reveal that decreased fitness can result from a catastrophic growth arrest caused by unexpected darkness in a small subset of cells with incorrect clock times corresponding to the subjective morning. We find that the clock generates rhythms in the instantaneous growth rate of the cell, and that time of darkness vulnerability coincides with the time of most rapid growth. Thus, the clock mediates a fundamental trade-off between growth and starvation tolerance in cycling environments. By measuring the response of the circadian rhythm to dark pulses of varying lengths, we constrain a mathematical model of a population’s fitness under arbitrary light/dark schedules. This model predicts that the circadian clock is only advantageous in highly regular cycling environments with frequencies sufficiently close to the natural frequency of the clock.

## Introduction

*Synechococcus elongatus* PCC 7942 (*S. elongatus*) is a photosynthetic, unicellular cyanobacterium that has been extensively used as a model system for the study of circadian rhythms (1, 2). Each cell contains a remarkably precise oscillator based on the *kai* genes (3). KaiA, KaiB, and KaiC work together to generate near-24 hour rhythms in the phosphorylation of the core clock protein KaiC, forming a biochemical oscillator that can be reconstituted *in vitro* (4, 5). In the cell, rhythmic changes in KaiC signal through histidine kinases to exert genome-wide control of transcription (6-8) and metabolism (9, 10).

Much is known about the behavior of this system under conditions of constant illumination, where robust cell-autonomous oscillations are easiest to observe (11-15). However, under constant conditions, *S. elongatus* can grow robustly even without a functioning clock (13, 16), leading us to suspect that the importance of the clock would be revealed by monitoring cellular physiology under conditions that fluctuate between light and dark. Indeed, previous work has shown that fitness defects occur in fluctuating environments with schedules that do not match the circadian clock period (1). Because environmental challenges may reveal heterogeneous behavior in a population, we designed a microscopy system that allows us to quantitatively measure clock state, growth rate, and cell division in individual cyanobacterial cells over several days in an environment that fluctuates between light and dark (Fig. 1, Movie S1). Using these single-cell measurements, we then develop a phenomenological model where growth rate and the probability of surviving the night are determined by the current clock state, which is itself updated following each light-dark transition. This model provides a framework to calculate the impact on organismal fitness from a circadian clock driven by an arbitrary fluctuating environment.

**Figure 1.**
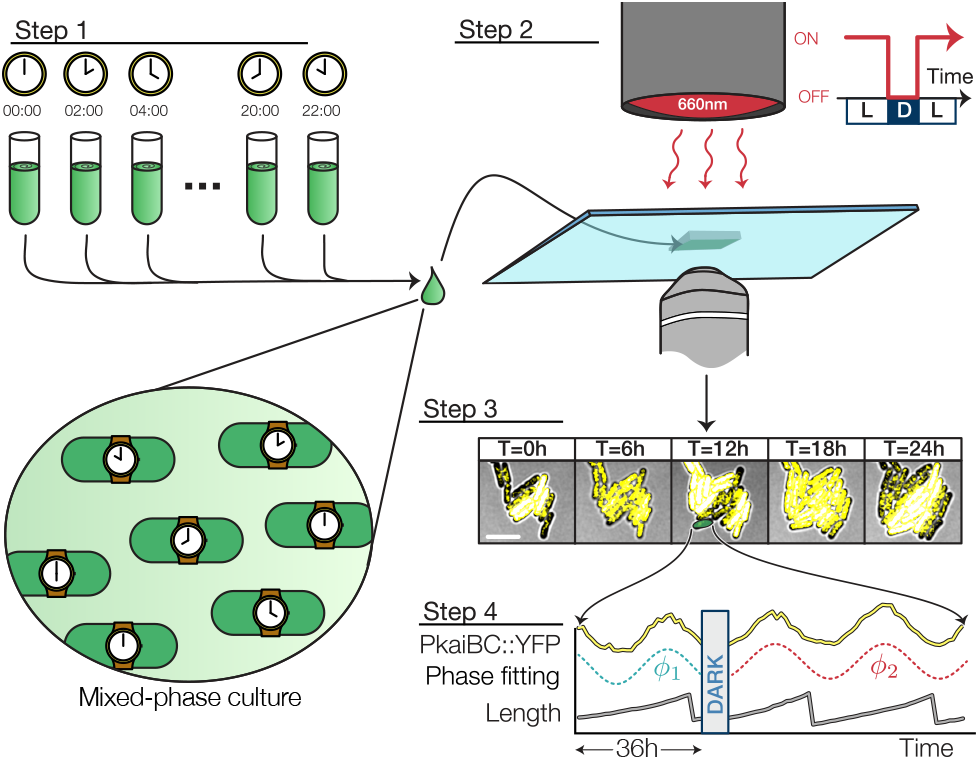
Experimental setup. 12 populations were entrained under staggered LD 12:12 regimes and combined into a single experiment. A multiplexed measurement of phase shift or growth rate modulation was achieved by exposing the mixed-phase population to a single pulse of darkness (scale bar = 5 µm). Fluorescence and brightfield micrographs recorded every hour were used to extract every cell’s physiological parameters (e.g. length, clock reporter).

**Figure 2.**
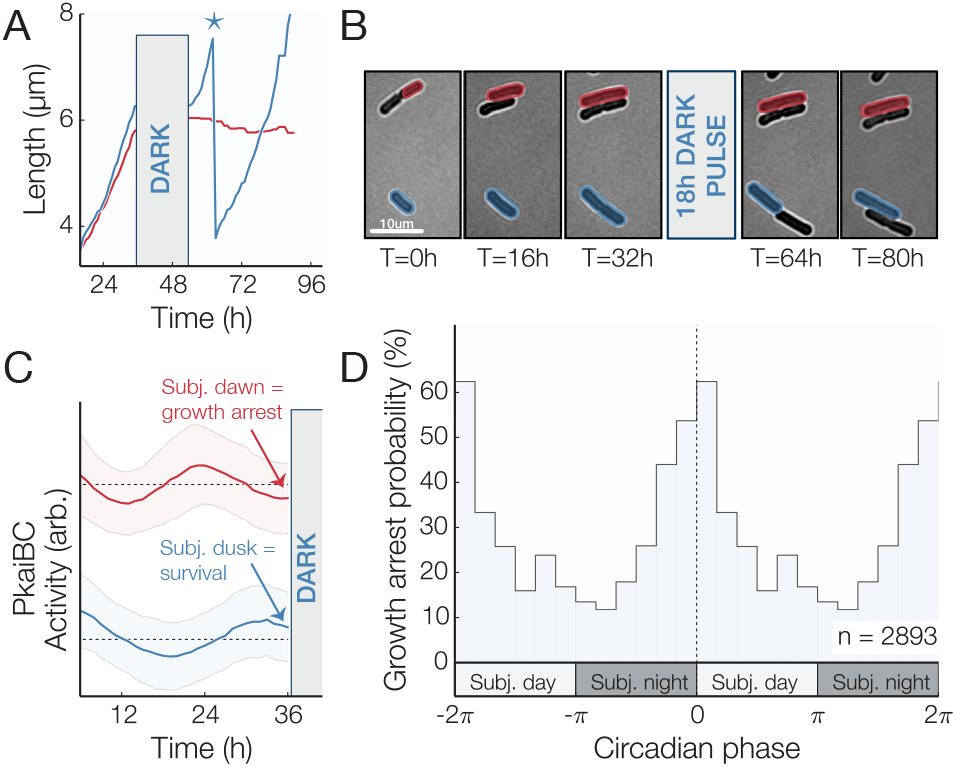
Clock-dependent growth arrest following unexpected darkness (A) Individual growth curves showing survival (blue) and growth arrest (red) of two neighboring cells (* = cell division). (B) Example of cells that entered a state of arrested growth following an 18h pulse of darkness. Cells remained dormant after >36h in constant light. (C) Average pre-darkness clock reporter time trajectory for surviving (blue) and arrested (red) populations. (D) Phase-dependent probability of growth arrest for cells grown under constant light conditions for 36h before being subjected to a 18h pulse of darkness (n=2983). The maximal growth arrest probability occurs when a dark pulse occurs near subjective dawn (transition between subjective night and day). Growth arrest probability is double plotted to illustrate its periodicity. See also Figure S1.

## Materials and Methods

### Cyanobacterial strains

The clock phase was tracked using a yfp-ssrA reporter strain WT/JRCS35 (MRC1006) that carried a PkaiBC::eyfp-ssrA fluorescence reporter. The JRCS35 plasmid integrated PkaiBC::eyfp-ssrA into NS2 (neutral site 2) with a kanamycin resistance cassette (12). To create the ∆*kaiBC* strain (MRC1009), the WT/JRCS35 strain was transformed with plasmid MR0091, replacing the endogenous *kaiBC* locus (from the *kaiB* start codon to approximately 200 bp upstream of the *kaiC* stop codon) with a gentamicin resistance cassette. The KaiBC overexpression strain (MRC1010) was created by transforming the WT/JRCS35 strain with plasmid MR0095, integrating *kaiBC* under control of the IPTG-inducible trc promoter into NS1 (neutral site 1).

### Culture conditions

In all the experiments, cyanobacterial strains were grown in BG11 liquid medium supplemented with 20 mM HEPES (pH 8.0) at 30◦C. To create the mixed-phase population, 200uL of a cell culture grown under continuous illumination (LL) 75 µmol photons m^-2^s^-1^ were pipetted into each well of a black (opaque) 96-well plate. For the experiments that included either the *kaiBC*-null or *kaiBC*-overexpression strains, these strains were grown in separate wells from the wild-type cells within the same plate as to expose them to the same culture and illumination conditions. For the *kaiBC*-overexpression experiment, the media was supplemented with IPTG at a final concentration of 1 mM within the plate. A custom-made Arduino driven LED array was used to illuminate each well. Each output pin of the Arduino supplied 23 mA of current to 8 red LEDs and each pin corresponded to one column of the 96-well plate. The Arduino was programmed to generate 2 days of symmetric light/dark conditions (light conditions: 10 µmol photons m^-2^s^-1^ (23 mA); Dark conditions: 0 mA) preceded by at least 12h of continuous light conditions so that each population was subjected to 2 entrainment cycles. Light levels were maintained at ∼10 µmol photons m^-2^s^-1^ for an additional 24h before cells were collected for microscopy. Each culture well of the entrained 96-well plate was collected and combined into a single test tube. For experiments that included *kaiBC*-null or *kaiBC*-overexpression strains, the cells were combined in equal proportions, determined by OD750 measurements after entrainment. This provided a mixed population of wildtype and mutant cells within a single experiment, as in supplementary movies S3 and S4.

### Timelapse microscopy

The mixed-phase culture was diluted to an optical density OD750 = 0.1 using BG11 medium and 1µL of the cell solution was pipetted onto a glass-bottom 6 well plate (MatTek Inc.). A small (1mm X 1mm X 0.5mm) pad of BG11 + 2% low-melting point agarose (LMPA) was placed atop the cell suspension. 10 mL of liquid BG11 + 2% LMPA which had been cooled to 37◦C was then poured inside the well to cover the LMPA pad. For the *kaiBC*-overexpression experiment, the BG-11/agar mixture was supplemented with IPTG at a final concentration of 1 mM before pouring into the well. Once the LMPA solidified, the 6-well plate was then moved to a motorized microscope (IX71, Olympus) and fluorescence and brightfield images were recorded every 60 minutes. Control of the microscope was carried out using micromanager (17). Every 60 minutes, a motorized microscope stage (Prior) visited 24 pre-assigned locations containing at least 10 cells and bright-field (exposure: 100ms), chlorophyll (exposure: 200ms; excitation: 501 nm; emission: 590 nm) and YFP fluorescence (exposure: 2s; excitation: 501nm; emission: 550 nm) micrographs were then recorded using an EMCCD camera (Luca, Andor). The “simple-autofocus” routine provided by the Micro-manager suite used the chlorophyll autofluorescence of the population to identify the focal plane before each set of micrograph was recorded. A collimated LED light (Thorlabs; wavelength: 625 nm) was used to illuminate the cells throughout the experiment and a microcontroller (Arduino) controlled the output level of the LED light (Light conditions:~10 µmol photons m s (23 mA); Dark conditions: 0 mA).

### Single-cell analysis and phase information extraction

The outline of every cell in the brightfield image was traced using a watershed algorithm and the physiological properties of each cell (length and YFP fluorescence intensity) were recorded. The celltracker image processing (18) suite was then used to reconstruct the lineage history of each cell, assigning an age to each pole and computing the instantaneous elongation rate.

The complete lineage of every cell present at the onset of the dark pulse was reconstituted and the YFP signal of each lineage was then subjected to a Fourier transform. The (complex) factor multiplying the 24h frequency component (henceforth called c24) was computed using the last 36h of data leading to the dark pulse. The phase of the cell before the dark pulse (called cp_1_ in the main text) was found by extracting the angle of c_24_ using the arctan2 branching function – ie. ϕ = arctan2(imag(c_24_), real(c_24_)).

To find the phase of the clock after the dark pulse (ϕ_2_), the YFP-intensity of the “old-pole” lineage (ie. the lineage which inherited the oldest pole after each division) was extracted and the phase information was found by computing the angle of the c_24_ factor of the YFP signal. After the dark pulse, only the first 36h of data were considered (to ensure that c_24_ existed). Non-oscillatory cells (such as the *ΔkaiBC* and *kaiBC*-overexpression strains) were identified by monitoring intensity traces which varied by less than 30% over the duration of the experiment.

Since the production and maturation rate of YFP proteins have a finite timescale that is determined, among other factors, by the growth rate of the cells, the clock phases we report are shifted by 4h relative to the extracted peak YFP phase information to bring in more closely in line with the estimated peak transcriptional activity. This value is similar to other values reported in the literature (12)

### Growth arrest probability

“Arrested” cells were identified by tracking the cumulative increase in the total length of the cell after dark pulse. If the total length of a cell and its progeny increased by at least 33% after 24h the cell was scored as arrested. Because of the altered morphology of the *kaiBC* overexpression strain, we used an alternative test for these cells: if the relative elongation rate was less than 1% / hour over the 4 hours following the dark pulse, the cell was scored as arrested.

### Elongation rate measurements

The elongation rate was computed from 6 experimental replicates of mixed-phase populations grown under constant light conditions. Cells were grown for a total of 36h under constant light conditions and the last 12h (T=24–36h) were used to compute the elongation rate. The instantaneous elongation rate was found by computing the relative increase in cell size between two consecutive frames. Specifically,

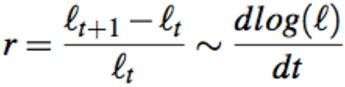

The growth rate *r*(*t*) was then binned according to the cell’s circadian phase and averaged over a 1h window. In Fig. 3B, a 3-pt moving average was used to smooth the data.

**Figure 3.**
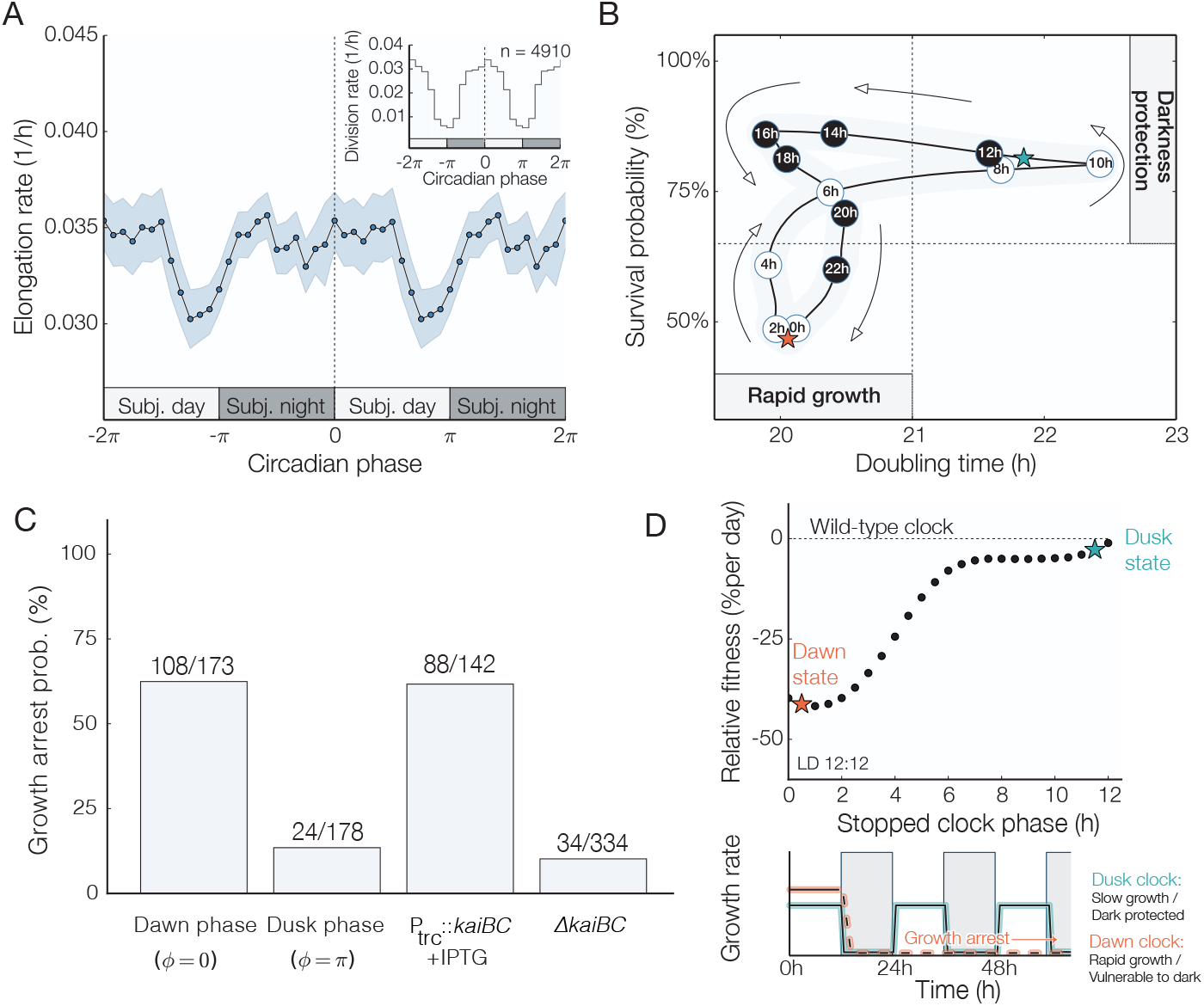
Clock dependent fitness trade-offs. (A) Elongation rate measurements (mean ±3×s.e.m) for cells grown under constant light conditions display a transient decrease at subjective dusk. Inset: Clock-dependent division rates computed from 4910 individual division events. Elongation and division rate measurements are double plotted to illustrate their periodicity. (B) Phenomenological model of the time-evolution of the circadian fitness trade-off. The subjective circadian time is inscribed inside each datapoint to show how a cell’s phenotype cycles between rapid growth and a starvation-protected state. The properties of the dusk and dawn phenotypes is marked with a cyan and orange star, respectively. (C) Growth arrest probabilities for wildtype and *kaiBC* mutant cells following an 18 h pulse of darkness. (D) Comparison between a clock stopped at dusk (cyan) or a dawn (orange). Since no phenotype exhibits superior fitness at all times, the performance of stopped-clock daytime strategies is lower than a circadian phenotype under circadian (LD 12:12) environments. Physiological states corresponding to subjective night (circadian times between 12-18h) were excluded from this analysis. Bottom: Cells in a dusk-like phenotypic state grow more slowly but are protected against the dark. Cells in a dawn-like phenotypic state grow more rapidly but are vulnerable to darkness. See also Figure S2.

### Robustness and phase resetting

The resetting index and the D5 phase shift of Fig. S3B were extracted from the 5h and 9h phase shift experiments (Figs. 3E and G). Graphically, the D5 phase shift index measured the average distance between the ϕ_1_ = ϕ_2_ dashed blue line in Fig. 4E and the resetting index measured the distance individual datapoints and the ϕ_2_ = π dashed red line in Fig. 4G. Data was binned over a 1h window and noise was smoothed out using a 3-pt moving average.

**Figure 4.**
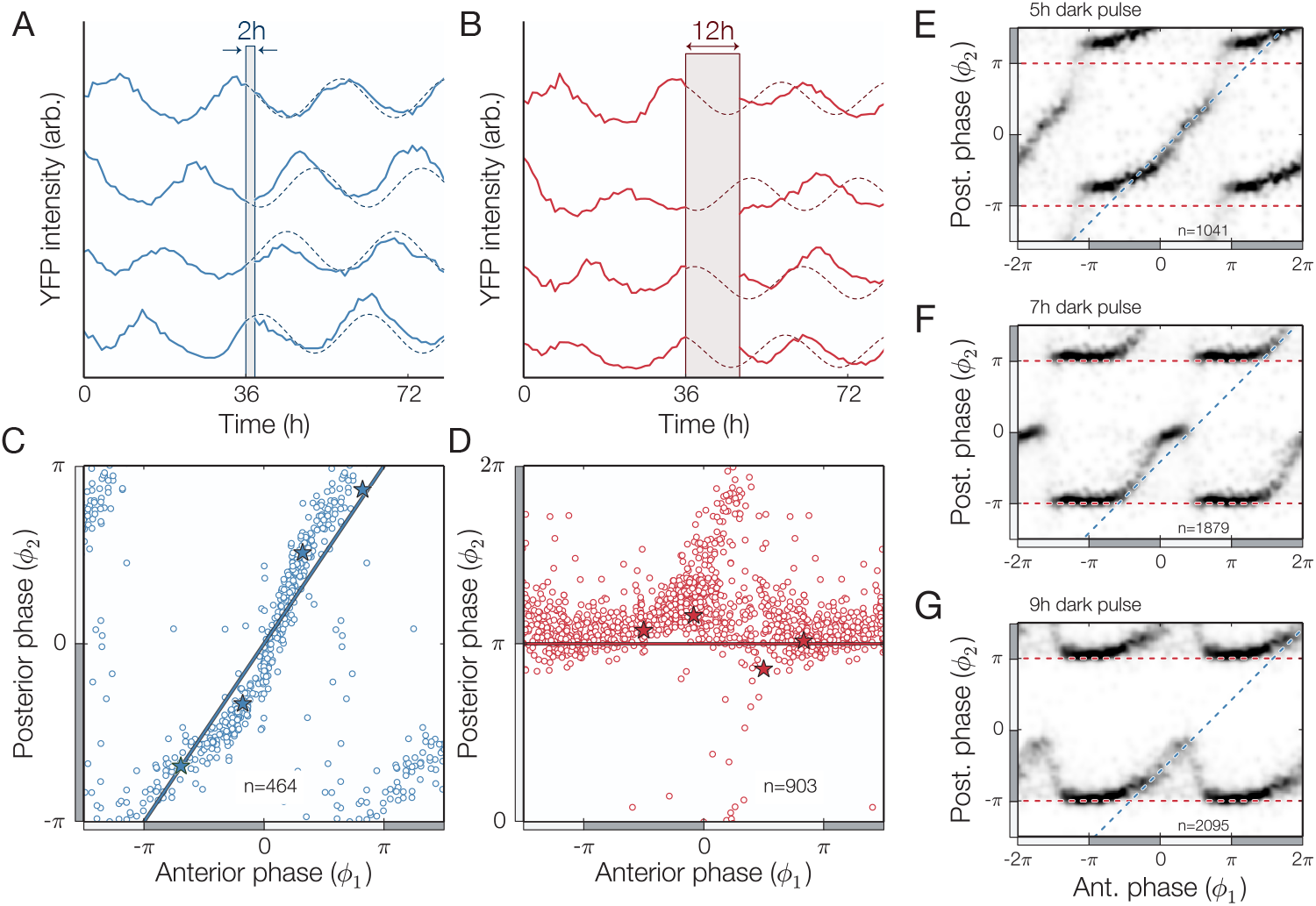
Single cell clock response to dark pulse perturbations. (A-B) Individual traces (Pk_a_iBC::eyfp-ssrA) showing phase-shifts caused by weak (2h dark pulse) and strong (12h dark pulse) perturbations. (C-D) Phase resetting of individual cells grown under constant light conditions for 36h before being subjected to a 2h (blue) or 12h (red) pulse of darkness, with specific examples from panels A-B are highlighted (*). Two distinct resetting behaviors are observed: a robust response (blue line in panel C) or a full phase reset (red line in panel D). (E-G) Density plot showing the phase response to 5h, 7h, and 9h dark pulses, with resetting that switches between no response and full-reset (blue and red dashed lines, respectively). Cells were grown under constant light conditions for 36h prior to each dark pulse. In all panels, the location of subjective day (night) is marked with a light (dark) gray bar. See also Figure S3.

### Fitness advantage measurement

To measure the performance of various clock periods under a sustained LD 12:12 schedule, it was necessary to compute 1) the probability µ that a cell would enter a state of growth arrest following the 12h dark pulse and 2) the number of cell doublings that happened during the 12h of light. µ was determined using 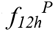 to find the phase of the cell at “dusk” to identify the survival probability at that phase using Fig. 2D (that is, it was assumed for the sake of simplicity that growth arrest occurs at the same rate for 12h and 18h nights). The number of cell doublings was found using 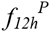 to identify the phase at the beginning of the day. This phase over the whole day was then used to compute the average elongation rate 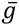. The historical fitness (19) was used to quantify the fitness of the population at a given period.

The historical fitness was given by

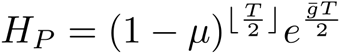

for a simulation that lasted for a time *T* for a given clock period *P*. The values of *H_P_* were plotted relative to *H*_24*h*_.

## Results and Discussion

### A subset of cells with misaligned clocks do not survive the night

Some photosynthetic organisms that rely exclusively on light for growth are known to halt DNA synthesis and enter a dormant state (20), or even die (21), in the absence of light. Since control of gene expression in the dark and consumption of energy metabolites are both under active control of the circadian clock in cyanobacteria (9, 10, 22), we hypothesized that unanticipated nightfall at clock times when energy reserves are low and metabolic rates are high could have deleterious effects. To test this hypothesis, we exposed a uniformly distributed (Fig. S1) mixed-phase population to a period of darkness corresponding to a long night (18h). Surprisingly, we found that a subset of cells experienced a catastrophic growth arrest after the simulated night: growth of these cells ceased and did not resume even after 36 hours of subsequent light exposure (Figs. 2A-B, Movie S2). To determine whether the ability to tolerate darkness-induced starvation is influenced by the circadian clock, we assigned a clock time to each cell by measuring rhythms in a fluorescent reporter of clock gene expression prior to the dark pulse. We found that the fraction of cells that failed to resume growth was strongly enriched for cells with clock states corresponding to the early day, when nightfall is not anticipated. Indeed the probability of dark-induced growth arrest oscillates with clock time, reaching a minimum at subjective dusk when nightfall is expected to occur (Figs. 2C-D). Thus, the ability of individual *S. elongatus* cell to tolerate prolonged starvation is clock-dependent, with cells displaying enhanced starvation tolerance when the onset of darkness coincides with subjective dusk.

### The clock allows rapid growth early in the day

In many microbes, stress tolerance is generally anticorrelated with growth rate (23). A classic example is the bacterial stringent response to amino acid starvation: mutants that cannot mount the stringent response can grow faster than the wildtype as nutrients are being depleted, but these mutants cannot survive conditions of prolonged starvation (24, 25). We therefore asked whether the rhythmic dark tolerance we observed in cyanobacteria is similarly linked to a change in growth rate during the circadian cycle.

By tracking morphological changes in single cells, we assigned an instantaneous growth rate to each cell and identified cell division events. We found that subjective dusk, the time when starvation resistance is highest, is also a time of slowed biomass incorporation (Fig. 3A). This time of slowed growth approximately coincides with the previously reported (26, 27) clock-controlled inhibition of cell division (Fig. 3A, inset). This reduction in cell growth and division is anticorrelated in time with the vulnerability of cells to darkness, suggesting the existence of a fundamental trade-off between the capacity for rapid growth and the ability to tolerate starvation (Fig. 3B).

To determine if these changes in dark tolerance are indeed caused by signaling from the circadian clock, we repeated these experiments using cells with either the *kaiBC* genes deleted or overexpressed under the control of an IPTG-inducible promoter. Based on previous studies, we expect deletion of *kaiBC* to result in arrhythmic high expression of dusk-expressed genes and elevated glycogen levels, mimicking a dusk-like state (3, 10). Conversely, we expect overexpression of *kaiBC* to cause arrhythmia while repressing dusk genes (28).

Consistent with the expectation that their physiology is dusk-like, we find that *kaiBC*-null cells fail to efficiently undergo cytokinesis and some cells exhibit filamentous growth under the microscope (Movie S3). Further, the *kaiBC*-null mutant is quite dark-tolerant and shows a slightly lower elongation rate but a much higher survival rate, independent of the timing of darkness (Fig. 3C and Fig S2). In contrast, *kaiBC* overexpression makes cells highly vulnerable to a light-dark transition, and the majority of these cells do not survive our dark pulse treatment (Fig. 3C). When grown on the microscope, *kaiBC* overexpression leads to some cell death prior to the dark pulse, and to a surprising morphological defect where the cytoplasm appears to expand at a rate that is not properly balanced by elongation, causing cells to lose their rod-like shape (Movie S4).

We used these growth and survival data to calculate the expected fitness for a simulated population of cells with a constant growth rate and constant dark tolerance, according to the inverse relationship we observed for the wildtype. This calculation predicts that oscillating growth outperforms all fixed daytime growth strategies in 12h:12h light-dark cycles (Fig. 3D). Recent theoretical work suggests that organisms typically optimize evolutionary trade-offs by adopting a compromise phenotype that interpolates between “archetypes” that represent the extreme demands on the system (29). Our findings represent a dynamical version of this phenomenon where cyanobacteria are able to achieve higher fitness by cycling between incompatible states of growth and starvation protection.

### Response of single cells to dark pulses is all-or-none

Having characterized the impact of a pulse of prolonged darkness on clock-dependent growth, we sought to determine how the clock state in single cells is reset by pulses of darkness in order to build a model describing how cyanobacterial cells grow in arbitrary fluctuating environments. External cues, such as dark pulses, cause the cell to reset its phase in an attempt to bring the clock into alignment with the environment (30, 31). This phenomenon has been studied using bulk cultures of cyanobacteria (10, 32, 33), but such population-wide measurements may mask important features such as loss of coherence and amplitude attenuation because they are based on signals that represent the average of the oscillations coming from many independent cells.

We thus exposed populations of hundreds of cells in a spectrum of clock states to dark pulses of varying lengths and determined the clock phase both before (ϕ_1_) and after (ϕ_2_) the dark pulse. When the dark pulse was brief (2 hours) cells at all clock times were largely resistant to the perturbation, and we observed only small phase changes (Fig. 4A, C). In contrast, long dark pulses corresponding to the length of the night (12 hours) were capable of effecting nearly a full reset where most cells were synchronized to the onset of darkness (Fig. 4B, D).

Surprisingly, dark pulses of intermediate length produced a discontinuous combination of these responses. When the clock time was far from subjective dusk, the response of the system to a dark pulse was very small (Fig. 4E-G, data near the blue 1:1 lines). In a critical range of clock times, however, the response changed abruptly so that the system strongly synchronizes to the onset of darkness. The range of times when a full reset occurs grows as the dark pulse becomes longer (Fig. 4E-G, data near the ϕ_2_=π red lines). Biochemical studies of the Kai proteins have implicated changes in levels of metabolites during the night, such as the ATP/ADP ratio, with the ability of the circadian clock to reset its phase (30, 34). The abrupt change in resetting behavior we observe here for intermediate-length dark pulses may in part be caused by a timescale associated with depletion of key metabolites in the cell.

This abrupt change from insensitivity to strong sensitivity as clock time progresses may represent an optimal strategy for dealing with unexpected fluctuations in the environment, as it can ignore short dark pulses at times of day when they are unlikely (Fig. S3B). This behavior would be difficult to detect without high-resolution single-cell measurements, because averaging over many cells with slightly different phases would tend to blur out the true sharpness of the response (Fig. S3C).

### Mathematical model of fitness in fluctuating environments

We combined our measurements of clock-dependent growth and darkness-induced growth arrest (Fig. 5A) with darkness-induced clock resetting to build a mathematical model of cyanobacteria growing under arbitrary schedules of light and dark. We first asked how successfully the circadian clock could synchronize to a 24-hour day if the clock period were altered. In particular, the phase resetting information obtained in Fig. 4E-G was used to generate the mapping for different clock periods. The relationship between ϕ*_i_* and ϕ*_i+1_* was given by the recurrence relation 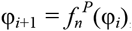, where the map 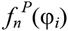 describes the phase at dusk following a dark pulse of duration *n* subjected to a clock of period *P*. The precise shape of 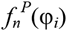 was found by interpolating the phase resetting curves by assuming that the effect of a dark pulse of the phase scales with the period of the clock (for instance, a 5h dark pulse would have the same effect on a 24h clock that a 10h dark pulse would have on a 48h clock). The shift between the phase before (ϕ*_i_*) and after (ϕ*_i+1_*) was used to construct an expression for 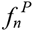 that accurately captures the features of the 5h, 7h, 9h, and 12h dark pulse measurements. In particular, we used the following expression to model 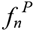:

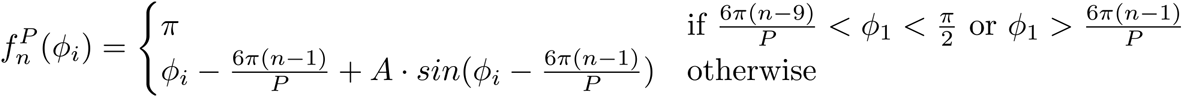

where *A* was given by min(n/P, π/4). Plots of this function for various dark pulse lengths are shown in Fig. S4.

**Figure 5.**
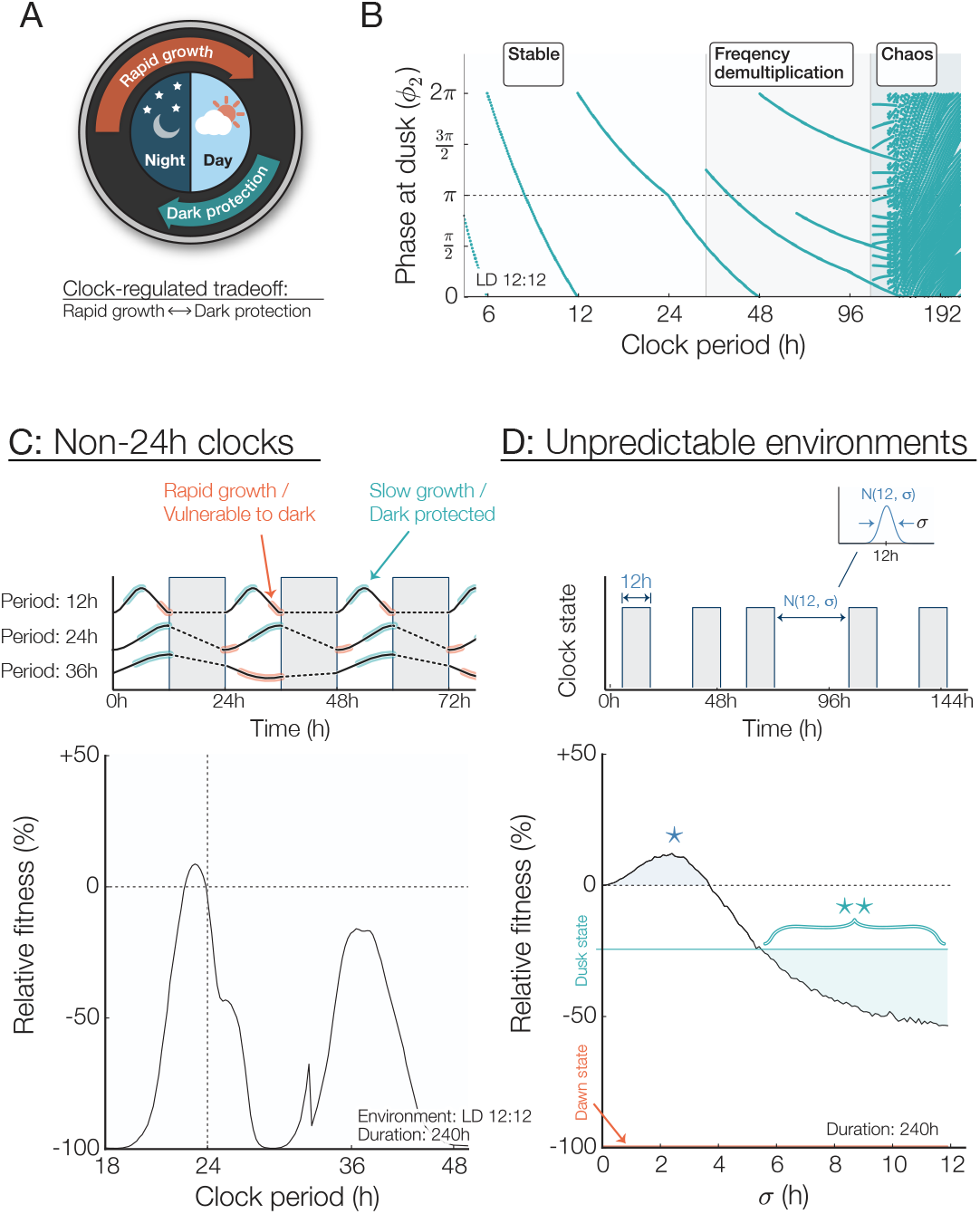
Mathematical model of clock-regulated growth under noisy and period mismatched schedules. (A) Circadian trade-off between rapid growth and stress protection. Each phenotype occupies a specific region of the circadian cycle. (B) Map showing phase entrainment, or lack thereof, under LD 12:12 for different clock periods. A perfectly entrained clock would result in a phase at dusk equal to π. Clock periods longer than 40h result in faulty phase tracking caused by frequency demultiplication and those longer than 100h lead to chaos. (C) Top: Effect of a non-circadian clock period. Depending on the clock period, phase entrainment can either be stable but inaccurate (clock period: 12h), stable and accurate (clock period: 24h), or unstable and inaccurate (clock period: 36h). Bottom: Relative fitness of various clock mutant subjected to a LD 12:12 circadian environment for 240h. Fitness advantage is the greatest when the clock and the environment are in constructive resonance (i.e. 24h and 36h clocks, which lead to a correct nightfall prediction) and the lowest for destructive resonance (i.e. 18h, 30h, 48h clocks, which results in nightfall occurring during subjective morning). (D) Top: Fitness under 12h nights but variable day lengths. Day durations are normally distributed with a 12h average and a variance σ. Bottom: The presence of a low level of noise in the day length distribution increases fitness for wild-type clocks (region highlighted with a *). As the day length becomes more unpredictable (σ > 5h), an “always-protected” dusk state is more beneficial than a circadian phenotype (** region). Fitness is the average of 10,000 simulations.

Using this expression for 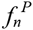 we interpolated from our measured dark pulse response data to find a stable recurrence relation corresponding to clock entrainment by plotting the value(s) of ϕ*_i_* which converged for *i* ≫ 1 for a 12h dark pulse each clock period. (Fig. 5B). When the clock period is less than 40h, the model predicts that the clock will stably entrain to the environment, but a period mismatch results in an entrained phase that is generally incorrect (i.e. subjective dusk does not fall near actual nightfall). For longer periods, entrainment can occur to subharmonics of the environmental period; for still longer periods, chaotic dynamics can occur.

We next investigated how circadian misalignment would affect the long-term fitness of a population of cells in the model by using the growth and survival functions measured in Figs. 1-2 to extract the fitness of a single cells under a given light/dark regime. By assuming that growth occurs only under light conditions and growth arrest probability is dependent on the phase at the onset of darkness, we calculated growth and survival of cells with a range of clock periods and find that the model predicts that the long-term fitness is highest when the clock period is near 24 hours.

This result indicates that the measured effects on growth and darkness tolerance may be sufficient selective pressures to explain the precision of the circadian clock. However, when the clock period is far from 24 h, large fitness costs can occur because the clock synchronizes inappropriately so that the morning clock state occurs at nightfall every day (Fig. 5C). These model results show similar trends to those previously reported from competition experiments: a long period (30 h) clock mutant is severely disadvantaged relative to wildtype in 12:12 LD cycles, while a short period (23 h) mutant is more mildly affected (1).

How does the circadian rhythm optimize the fitness of a cell? In our model, cells must grow slowly near nightfall to avoid the possibility of metabolic catastrophe when darkness falls. The clock enforces this growth slowdown late in the day while allowing cells to grow rapidly in the morning. This suggests that the advantages conferred by a circadian clock are a result of a finely tuned match with the temporal structure in the environment. Such a strategy might become detrimental in unnatural conditions where the environment cycles irregularly between light and dark. To test this hypothesis, we simulated cyanobacteria growing in days with random variation by selecting the duration of each light period from a normal distribution with a mean of 12 h. The model predicts that when the variability in the light-dark schedule exceeds σ=5h, an arrhythmic, slow-growing strategy similar to deletion of the *kaiBC* genes becomes more successful than the wildtype (Fig. 5C). Surprisingly, our model predicts that wildtype cells may grow faster and achieve a higher fitness in the presence of some timing variability (Fig. 5C, σ < 3.75h) in the duration of the day.

## Conclusions

Despite the ubiquity of circadian clocks, it has remained challenging to pinpoint the benefits of rhythmic physiology (35). Our ability to detect the costs and benefits of clock function at the single cell level provides a framework to answer these questions. We found that considerable fitness penalties result from the failure of cells to correctly predict the withdrawal of energy associated with nightfall. We thus conclude that a major function of the cyanobacterial circadian clock is to provide a safeguard against darkness-induced starvation, giving the cell permission to grow rapidly early in the day.

A possible explanation for the failure of a subset of cells to survive the night is that these cells might have been unable to properly manage their energy consumption over the length of the night. Analysis of microarray expression data of *S. elongatus* (8) provides some mechanistic insight into the origin of clock-dependent starvation tolerance (Supplementary Table I). Expression of genes involved in photosynthesis and the biosynthesis of essential compounds peaks at clock times corresponding to the early morning, suggesting that the clock tunes metabolism to allow rapid growth early in the day. On the other hand, genes involved in DNA replication, DNA repair, and metabolism under nutrient limitation, peak late in the day, suggesting clock-dependent activation of mechanisms needed to tolerate nightfall. Furthermore, we previously found that the clock controls storage and consumption of energy storage metabolites, with reserves of glycogen at their lowest near the beginning of the day (10). Coupled with gene expression data, our results suggest that proper temporal regulation of energy storage and circadian regulation of growth and division in anticipation of dusk may play a critical role in allowing the cell to survive the night. Interestingly, cells early in the subjective night (i.e. circadian time between 14 – 18) are able to both grow rapidly *and* survive prolonged darkness. We suggest that this phenomenon is an example of phenotypic memory (18), wherein previous adaptations—in this case production of glycogen and other stress-protective factors during the slow growth portion of the cycle—are retained to confer an adaptive phenotype after the source of stress or stimulus has been removed.

Our results suggest an alternative to the hypothesis that circadian rhythms evolved primarily as a means to anticipate and avoid light-induced photodamage, i.e. a “flight-from-light scenario. If our experimental conditions approximate the challenges faced by the ancient ancestors of modern cyanobacteria, the daily threat posed by an extended time of resource limitation during the night may have been a major selective pressure on primordial clock systems. That is, a key function of the circadian clock is to direct preparations ahead of nightfall, i.e. a “prepare-for-night scenario.

Our dark pulse experiments show that the circadian clock has robustness properties that allow it to track the 24-hour cycle in the environment even in the face of random fluctuations. The flipside of this robustness is a remarkable fragility in environments that fail to have a 24-hour periodicity. Although such environments are unlikely to occur in nature, poor performance of clocks in these conditions may have relevance to other organisms that display a clock control of cellular divisions (36, 37) and to the irregular work schedules and patterns of light exposure typical of modern society.

## Acknowledgements

We thank Alexander van Oudenaarden (Hubrecht Institute, Utrecht) for the generous gifts of plasmids and Gopal Pattanayak for assistance with molecular biology. We thank Michael Glotzer, Joe Markson and members of the M.J.R. lab for comments on the manuscript. The work was supported by a Burroughs-Wellcome Career Award at the Scientific Interface (M.J.R.), the Chicago Fellows program (G.L.), NIH training grant T32-GM007281 (J.C.), and by an NIH R01 award (GM107369-01). Author contributions: G.L. and M.J.R. designed the study and wrote the paper. G.L. and J.C. carried out the experiments, and G.L. analyzed the data. The authors declare no conflicts of interest.

